# Conformational diversity and protein-protein interface clusters help drug repurposing in Ras/Raf/MEK/ERK signaling pathway

**DOI:** 10.1101/2023.08.03.551801

**Authors:** Ahenk Zeynep Sayin, Zeynep Abali, Simge Senyuz, Fatma Cankara, Attila Gursoy, Ozlem Keskin

**Affiliations:** Department of Chemical and Biological Engineering, Koc University, Istanbul, 34450, Turkey; Department of Computational Sciences and Engineering, Koc University, Istanbul, 34450, Turkey; Department of Computer Engineering, Koc University, Istanbul, 34450, Turkey

**Author notes:** Corresponding author: Ozlem Keskin College of Engineering, Chemical and Biological Engineering Rumeli Feneri Yolu Sariyer, 34450, Istanbul, Turkey Voice. **Abbreviations and symbols:** CDK4, Cyclin-dependent kinase 4; CDK6, Cyclin-dependent kinase 6; CDKN2D, Cyclin-dependent kinase inhibitor 2D; EGF, Epidermal growth factor; EGFR, Epidermal growth factor receptor; ERK, Extracellular signal-regulated kinase; FDA, Food and Drug Administration; HIV, Human immunodeficiency virus; HTR4, Human serotonin receptor 4; MAPK, Mitogen-activated protein kinase; MEK, Mitogen-activated protein kinase kinase; mTOR, Mammalian target of rapamycin; PDB, Protein Data Bank; PI3K, Phosphatidylinositol-3 kinase; PPI, Protein-protein interaction.

**Keywords:** Ras signaling pathway, protein-protein interactions, protein-protein interfaces, alternative conformations, drug repurposing

## Abstract

Ras/Raf/MEK/ERK signaling pathway regulates cell growth, division, and differentiation. In this work, we focus on drug repurposing in the Ras/Raf/MEK/ERK signaling pathway, considering structural similarities of protein-protein interfaces. The complexes in this pathway are extracted from literature and interfaces formed by physically interacting proteins are found via PRISM (PRotein Interaction by Structural Matching) if not available in Protein Data Bank. As a result, the structural coverage of these interactions has been increased from 21% to 92% using PRISM. Multiple conformations of each protein are used to include protein dynamics and diversity. Next, we find FDA-approved drugs bound to additional structurally similar protein-protein interfaces. The results suggest that HIV protease inhibitors tipranavir, indinavir and saquinavir may bind to Epidermal Growth Factor Receptor (EGFR) and Receptor Tyrosine-Protein Kinase ErbB-3 (ERBB3/HER3) interface. Tipranavir and indinavir may also bind to EGFR and Receptor Tyrosine-Protein Kinase ErbB-2 (ERBB2/HER2) interface. Additionally, a drug used in Alzheimer’s disease (galantamine) and an antinauseant for cancer chemotherapy patients (granisetron) can bind to RAF proto-oncogene serine/threonine-protein kinase (RAF1) and Serine/threonine-protein kinase B-Raf (BRAF) interface. Hence, we propose an algorithm to find drugs to be potentially used for cancer. As a summary, we propose a new strategy of using a dataset of structurally similar protein-protein interface clusters rather than pockets in a systematic way.

**Significance statement:** This work focuses on drug repurposing in the Ras/Raf/MEK/ERK signaling pathway based on structural similarities of protein-protein interfaces. The Food and Drug Administration approved drugs bound to the protein-protein interfaces are proposed for the other interfaces using protein-protein interface clusters based on structural similarities. Moreover, the structural coverage of protein complexes of physical interactions in the pathway has been increased from 21% to 92% using multiple conformations of each protein to include protein dynamics.

## INTRODUCTION

Drug repurposing or drug repositioning, using a drug for an indication other than its original purpose, is an attractive option compared to the long and costly process of developing a new drug^1, 2^. The drug to be repurposed has already been studied for its safety and has extensive data on its pharmacokinetics. As a result, many stages of drug development can be omitted^3^. Some examples of successfully repurposed drugs are thalidomide and sildenafil. Thalidomide, an antiemetic drug for pregnant women that was subsequently proven to have teratogenic effects, has been repurposed to be used in leprosy and sildenafil, a drug originally developed for angina, has been used in erectile dysfunction^4^. Current drug repurposing cases typically follow a disease-centric approach but when disease-focused repurposing reaches its limits, target-centric and drug-centric repurposing relying on structural data will be crucial ^5^. Docking and virtual screening are some of the most common methods in computational drug repurposing for preliminary studies ^6^. Some of the structure-based virtual screening web servers for drug repurposing are ACID (using inverse docking approach ^7^), DRDOCK (combining docking and molecular dynamic simulations for a target protein ^8^) and MTiOpenScreen (using docking or blind docking ^9^).

Cell signaling is the transmission of an external signal to activate certain mechanisms in the cell^10^. Ras/Raf/MEK/ERK signaling pathway plays a role in transduction of a signal received from an extracellular receptor to the cell nucleus to regulate biological functions, including cell proliferation, differentiation, apoptosis and stress response ^11–14^. Dysregulation of this pathway is associated with diseases such as inflammation, developmental disorders, neurodegenerative disorders ^11, 15–17^ and is observed in approximately one-third of all human cancers ^18^. Consequently, various drugs targeting this pathway have been developed. Vemurafenib, dabrafenib and trametinib are some examples of MAPK inhibitors used in cancer therapy ^19^. Proteins in this pathway interact with other proteins and these interactions take place through protein-protein interfaces ^20^. Hence, protein-protein interfaces are critical targets for drugs to regulate abnormal protein-protein interactions (PPIs) in this pathway ^21, 22^. Disruption of a PPI by targeting the interface with a drug may interrupt transduction of a signal that promotes tumorigenesis, thereby being beneficial in cancer therapy ^23^. From our previous studies we know different proteins can form similar protein-protein interface architectures ^24–26^. Using similar interfaces, Engin et al. ^27^ proposed that drugs binding to an interface might also bind to another interface with a similar structure. In their case study, they showed that the drugs binding to the interface between CDK6 and CDKN2D also bind to the interface between CDK4 and CDKN2D, which has a similar interface, with comparable binding energies.

Here, our aim is to use a non-redundant protein-protein interface dataset that is clustered based on structural similarity for drug repurposing. We preferred studying protein-protein interfaces rather than the binding pockets because in some cases target proteins may lack binding pockets, limiting their druggability such as the case of RAS protein family ^28–30^. Moreover, molecular glue is a new concept that may be used to make the targets druggable that were once considered as undruggable and the protein-protein interfaces are perfect for this approach ^31^. In this study, we focused on protein-protein interfaces of Ras/Raf/MEK/ERK signaling pathway. We studied interfaces that are available in Protein Data Bank (PDB) ^32^ and used PRISM web server ^33, 34^, a prediction tool for protein-protein interactions at the structural level, to predict the interfaces between any two physically interacting proteins of Ras/Raf/MEK/ERK signaling pathway when there is no experimental data. Proteins are dynamic and the conformational space is diverse. Availability of different conformations is crucial to find the right one that fits to the drug molecule PDB is getting richer with many conformations for a single protein. Here, we used an ensemble of conformations rather than taking a single structure in the predictions (Figure 1). Considering alternative conformations of each protein, the number of successfully predicted interactions increases ^35^. We extracted drugs already bound to the interfaces as candidates of drug repurposing for target proteins with structurally similar interfaces. Finally, we performed docking to propose drugs to be repurposed and found literature evidence showing that the algorithm we used here can be promising to suggest new uses for already known drugs.

**Figure 1.**
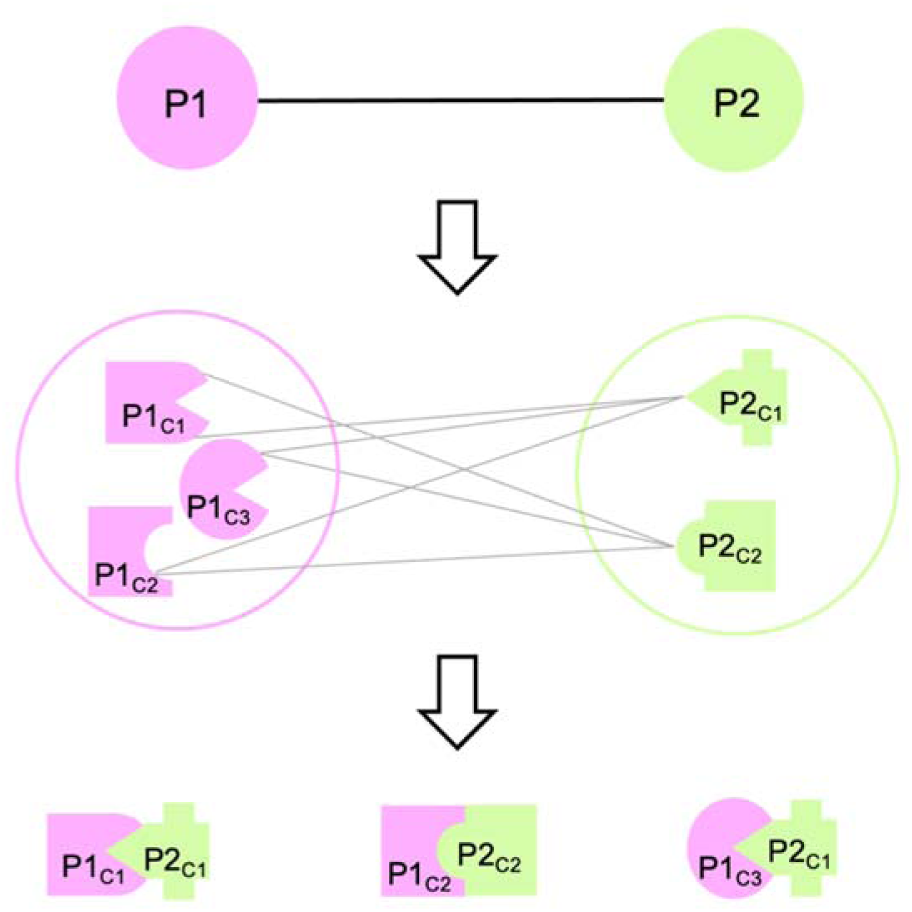
Protein-protein interactions with multiple conformations. Interacting proteins, Protein 1 (P1) and Protein 2 (P2) have three and two conformations respectively. Considering the multiple conformations of each protein, there are six possible interactions in theory (represented with grey edges) but in reality, only some of these interactions can be found (three of them in this case).

## RESULTS

The Ras/Raf/MEK/ERK signaling pathway is reconstructed by 16 proteins present in the KEGG database ^36–38^ under the EGF-EGFR-RAS-ERK signaling pathway and their top 10 interactors according to STRING database ^39^. All available structures of these 26 proteins in PDB ^32^ are grouped based on sequence and structural similarity. The representatives of the alternative conformation groups of these proteins can be found in Table S1. These conformations either correspond to the alternative conformations of the same region or they may correspond to different parts of a protein. The pathway proteins have 4.56 ± 5.20 conformations on the average. The structures of GAB2 in PDB have less than 30 amino acids and are eliminated in the grouping process. Its AlphaFold ^40, 41^ model is used in the following steps.

The network of Ras/Raf/MEK/ERK signaling pathway consists of 26 proteins and 72 interactions resulting in theory 2,564 possible interactions between alternative conformations. Interacting protein pairs considering alternative conformations are submitted to the PRISM web server ^33, 34^. PRISM simulations predicted 3,309 complexes for 66 of 72 interactions reported in STRING. These interactions can be seen in Figure 2. With the PRISM predictions, the structural coverage of protein-protein complexes formed by physically interacting proteins of this pathway has been increased from 15 to 66 out of 72 interactions. These results correspond to 999 of all 2,564 possible interactions among alternative conformations and through 630 unique protein interface templates (Text S1). The results involve some complexes for the same protein structures with different binding energy predictions. All the predictions are used in the next steps to avoid missing any new target.

**Figure 2.**
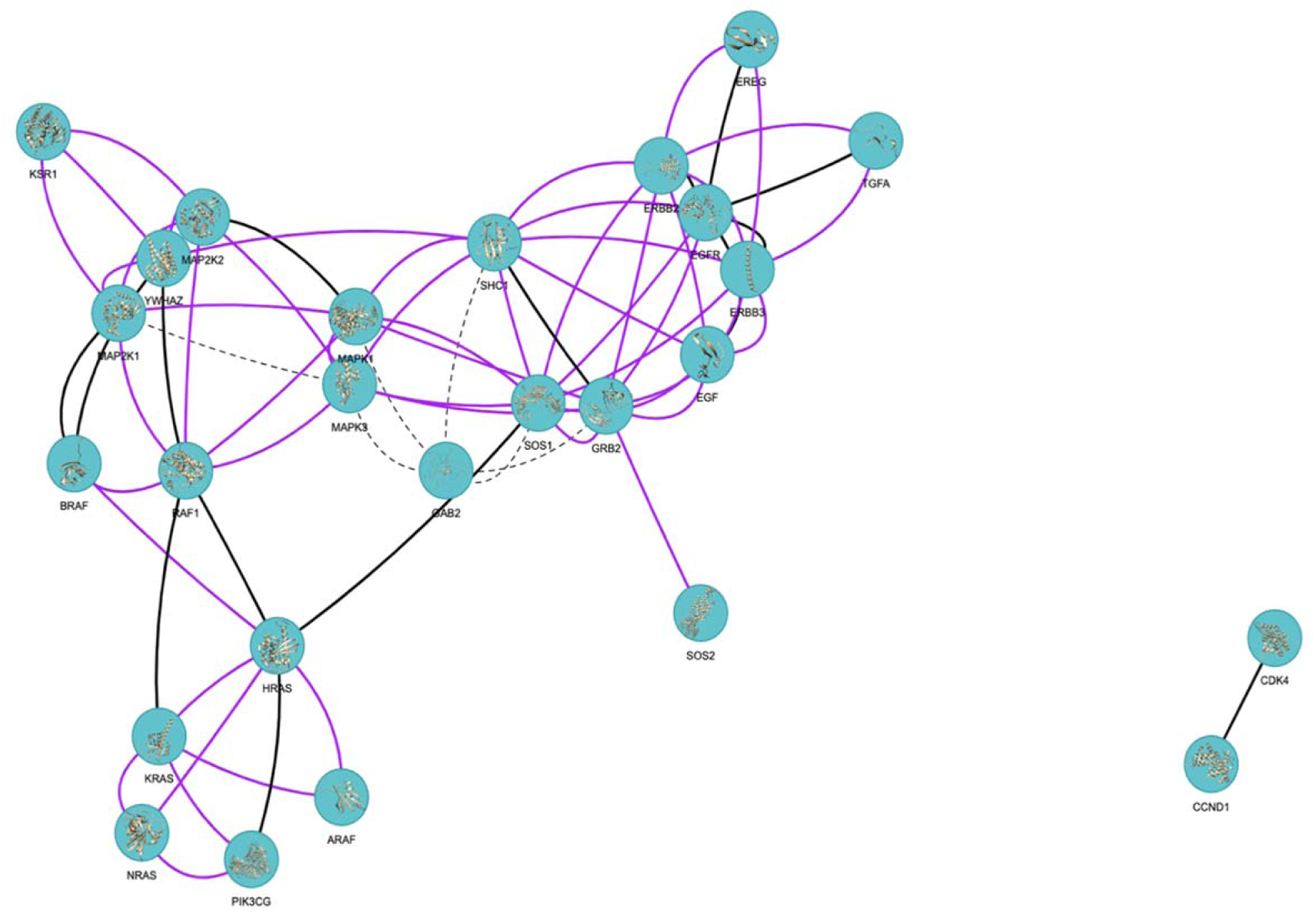
Protein-protein interaction network of Ras/Raf/MEK/ERK signaling pathway. Nodes represent proteins and proteins connected by edges represent the interaction between those proteins. If the edge is black, the complex of interacting proteins is available in PDB. If the edge is purple, the complex is not available in PDB but predicted by PRISM. If the edge is a dashed line, the complex is neither available in PDB, nor predicted by PRISM. ^42^

Additionally, there are 994 PDB structures, 521 of which have more than one chain, involving at least one of the 16 proteins in the EGF-EGFR-RAS-ERK signaling pathway. In total, there are 1,296 protein interfaces formed in these PDB entries. These interfaces are combined with the interfaces predicted by PRISM.

A structurally non-redundant dataset of protein-protein interface clusters (Dataset S3) ^43^ is used to find possible new drug-target pairs. A schematic representation of two scenarios is shown in Figure 3. The first one is “Repurposing To” where a drug bound to one of the interfaces in a protein-protein interface cluster may bind to an interface in the same cluster that belongs to the Ras/Raf/MEK/ERK pathway. In the second scenario of “Repurposing From”, a drug bound to a protein interface in Ras/Raf/MEK/ERK pathway may also bind to another protein interface that is in the same cluster, and the protein is not in the Ras/Raf/MEK/ERK pathway.

**Figure 3.**
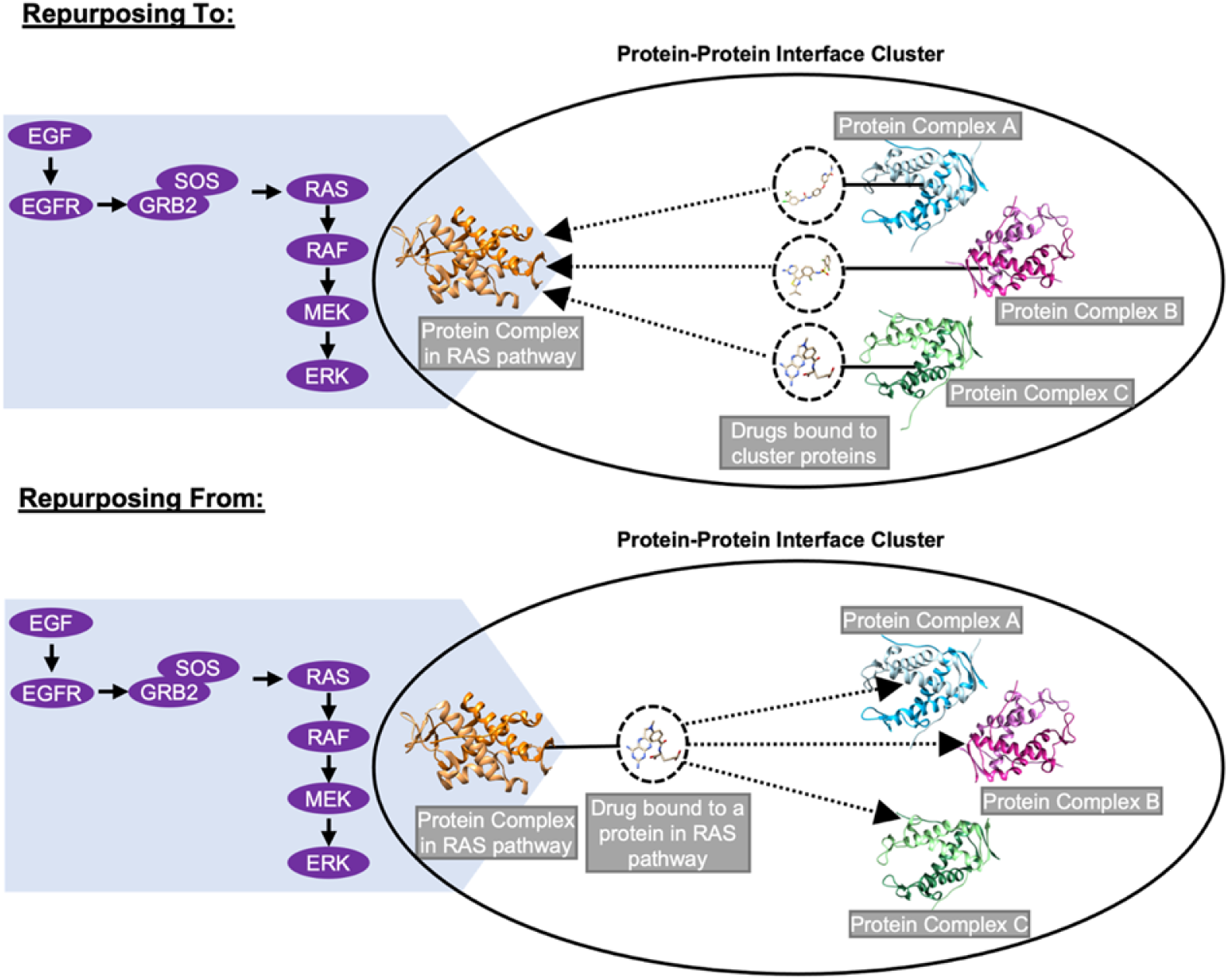
Identification of new drug-target pairs. Solid line represents a drug bound to an interface. Identified new drug-target pairs are represented with a dotted line with an arrow.

We filtered the clusters that contain all the template interfaces of the PRISM predictions and experimental PDB structures for the “Repurposing To” strategy. Templates of PRISM predictions are in 595 different clusters with a total number of 43 FDA-approved drugs bound to one of the protein interfaces in these clusters. Protein interfaces formed in PDB entries involving at least one of the pathway proteins are in 338 different clusters. The protein interfaces in these clusters have 12 unique FDA-approved drugs in total. With the approach of “Repurposing To”, there are 441 and 71 possible new drug-target pairs from PRISM results and PDB entries of pathway proteins respectively (Table S2).

Considering unique protein interfaces of PRISM predictions, there are 5 different FDA-approved drugs bound to 6 different protein interfaces. Whereas there are 8 protein interfaces with 3 different FDA approved drugs among the interfaces in PDB entries related to EGF-EGFR-RAS-ERK signaling pathway. The mentioned interfaces and drugs can be seen in Table S3. These protein interfaces are used for the “Repurposing From” strategy (see Methods for details).

All 6 protein interfaces with FDA-approved drugs among the PRISM predictions are in different clusters having 72 other protein interfaces in total. The 8 protein interfaces formed by the proteins related to our pathway are in 4 different clusters in which there are 60 other protein interfaces and there are no FDA-approved drugs bound to these interfaces. Consequently, with the approach explained as “Repurposing From”, we have 72 possible new drug-target pairs from PRISM predictions and 110 from protein interfaces in pathway PDB entries (Table S4).

We further performed docking for these drug-target pairs and the results are analyzed according to the binding free energy (Table S5). A previous study proposed an average binding energy of −7.75 ± 0.06 kcal/mol ^44^, accordingly new targets are presented in Table 1. These proteins contain both intracellular domains or extracellular domains and a result between intra- and extracellular regions is not biologically meaningful, therefore we eliminated such cases.

**Table 1.**
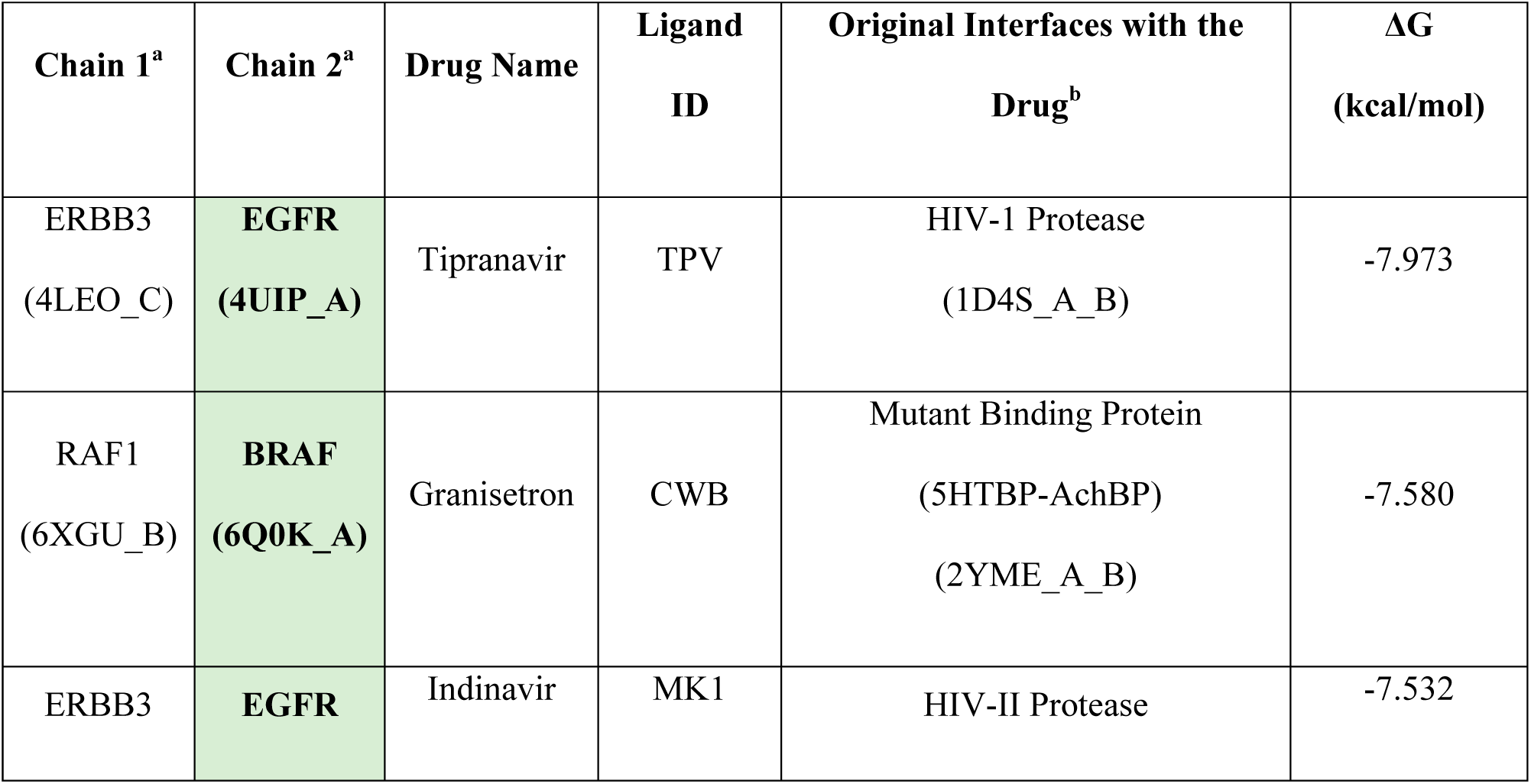
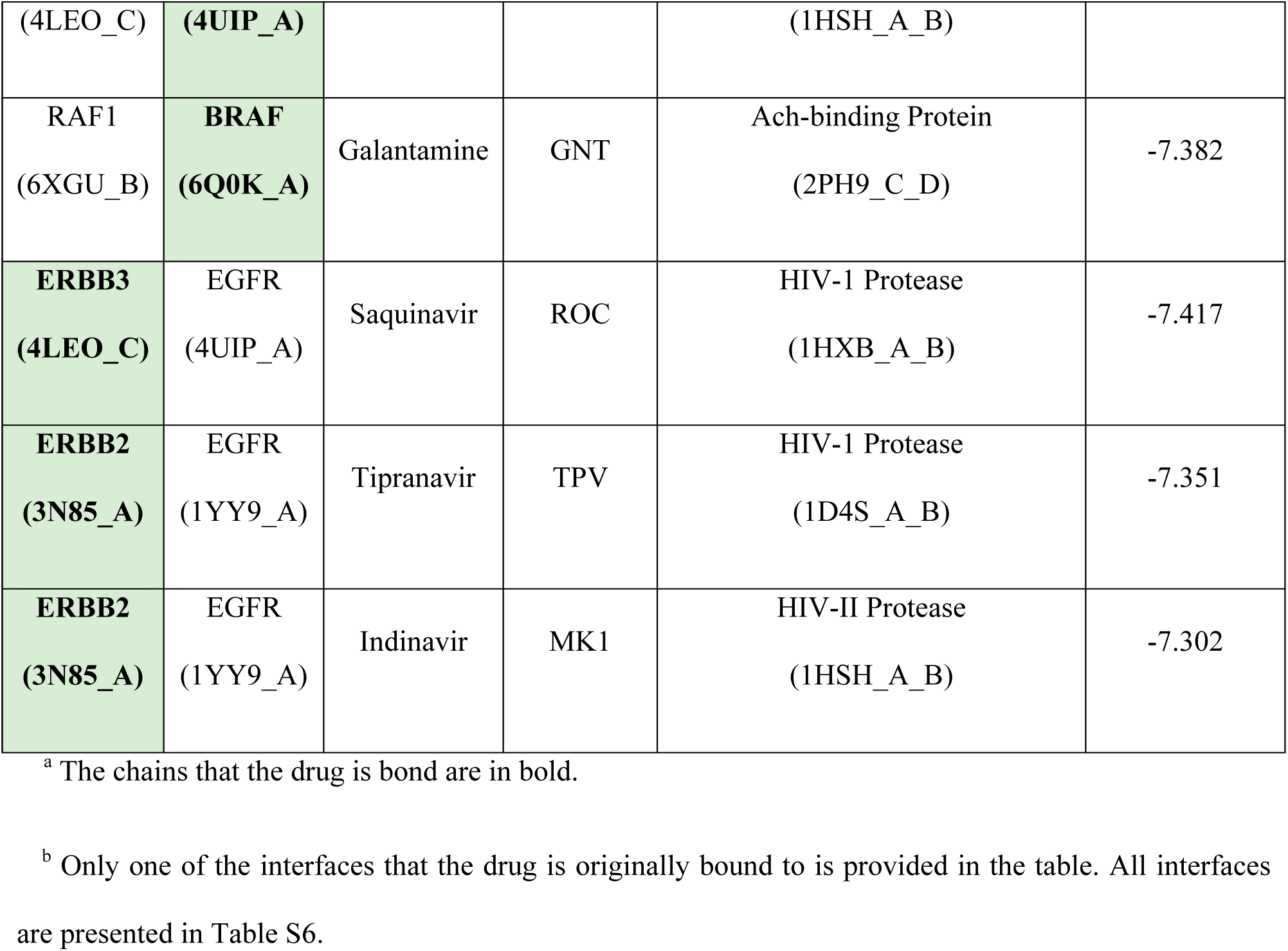
Proposed Drug Repurposing Candidates.

Table 1 presents the protein-protein interfaces proposed for drug repurposing (columns 1 & 2) and the protein complexes (column 5) with a drug bound to their interfaces that have structural similarity to the proposed interfaces. The structural similarity of BRAF-RAF1 interface with galantamine and the interface that galantamine is originally bound to is highlighted in Figure 4. In both cases the galantamine is bound to similar region of the interface. Moreover, the hydrogen bond between the galantamine and its original interface is also present between galantamine and BRAF-RAF1 interface.

**Figure 4.**
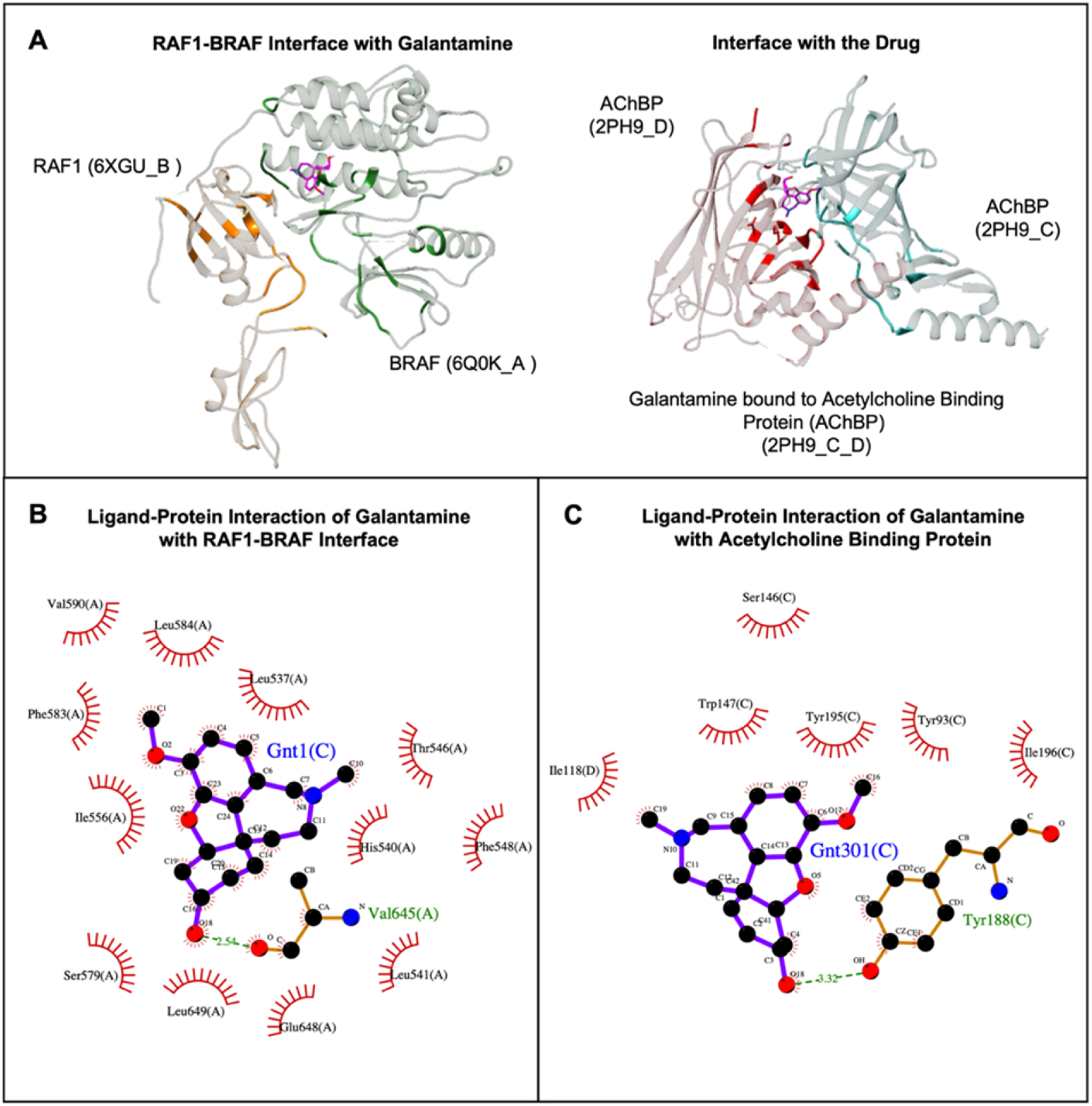
Similarity of the BRAF-RAF1 interface with galantamine and the interface that galantamine is originally bound to. (A) Structural similarity of BRAF-RAF1 interface and acetylcholine binding protein interface with galantamine (2PH9_C_D) is highlighted. (Molecular graphics and analyses performed with UCSF Chimera, developed by the Resource for Biocomputing, Visualization, and Informatics at the University of California, San Francisco, with support from NIH P41-GM103311.0) (B) Ligand-protein interaction diagram for BRAF-RAF1 interface and galantamine. There is a hydrogen bond between the galantamine and Val(645) of BRAF represented with green dashed line. (C) Ligand-protein interaction diagram for acetylcholine binding protein interface and galantamine. There is a hydrogen bond between the galantamine and Tyr(188) of acetylcholine binding protein (2PH9_C) represented with green dashed line ^45^.

Since mutations at the interface may alter the protein-protein interactions and the interaction with ligands, residues where cancer mutations are observed are extracted from the COSMIC database. Then, they are mapped to the interface residues. The cancer mutations located at the interfaces (of the protein-protein complexes in Table 1) can be seen in Figure 5. At the interface of the ERBB3-EGFR complex, ERBB3 has 6 residues and EGFR has 8 residues where cancer mutations are observed. There are 4 residues of RAF1 and 13 residues of BRAF related to cancer mutations at the interface of BRAF-RAF1 complex. Lastly, ERBB2-EGFR complex has 3 residues at the ERBB2 side of the interface and 8 residues at the EGFR side of the interface with cancer mutations. Mapped mutations for all the complexes used in this study are presented in Table S7 and Table S8.

**Figure 5.**
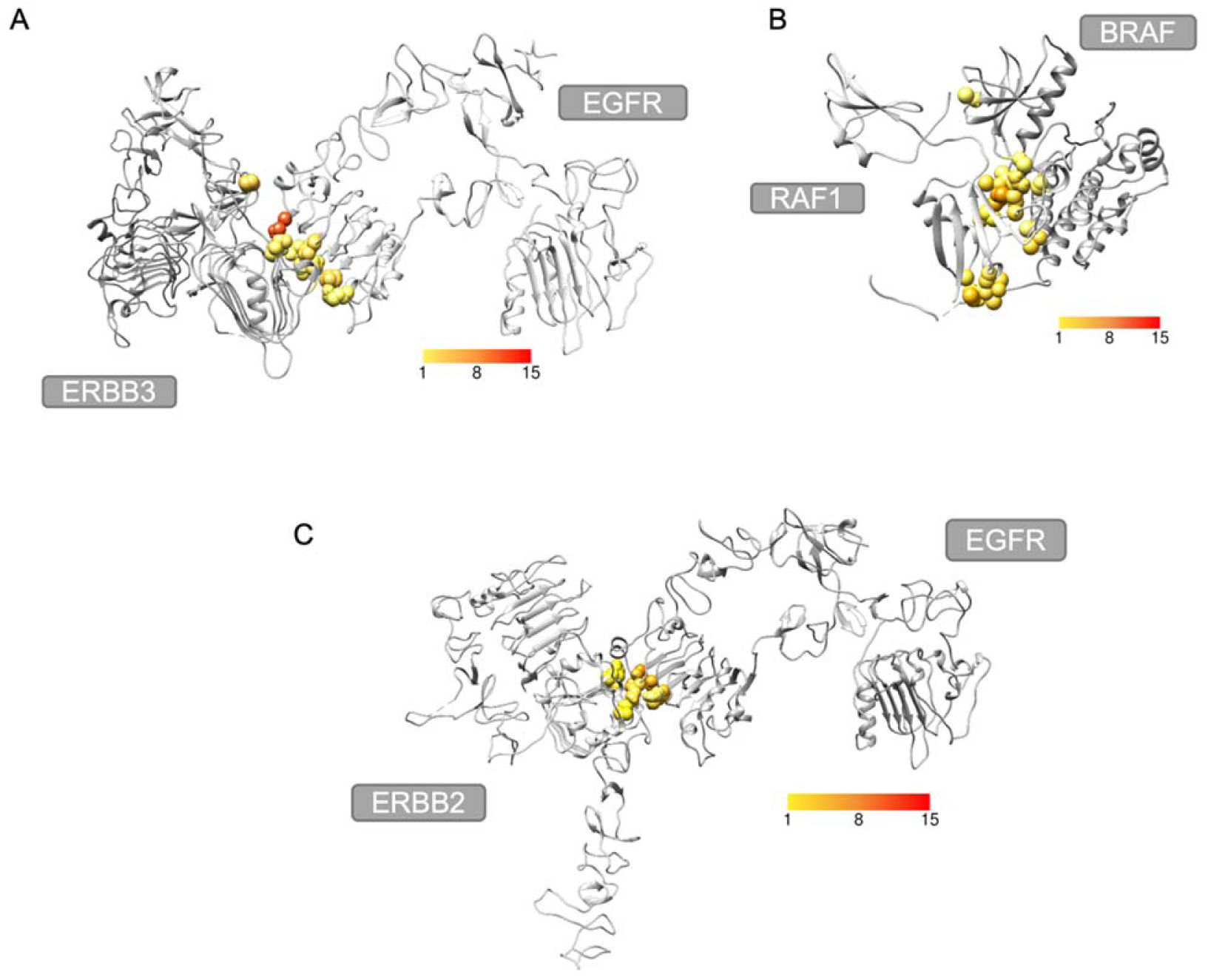
Cancer mutations at the interface of the protein-protein complexes (A) ERBB3-EGFR complex with cancer mutation residues colored according to frequency. (B) BRAF-RAF1 complex with cancer mutation residues colored according to frequency. (C) ERBB2-EGFR complex with cancer mutation residues colored according to frequency.

## DISCUSSION

Drug repurposing may adopt a ligand-based approach or target-based approach. Here, we used a new concept, the structural similarities of the protein-protein interfaces to propose new targets for the FDA approved drugs in Ras signaling pathway. This method required the 3-dimensional structures of the protein-protein complexes. However, only 21% of the physically interacting protein complexes were available in PDB for our network. Using PRISM, a template-based structural prediction, structural coverage of the network is increased to 92%. Here, conformational diversity of proteins are integrated by using multiple conformations of the pathway proteins. For instance, if just one structure (PDB ID:3KSY Chain ID:A) of SOS1 had been used, protein-protein complex structures for only 60% of the listed interactions would have been found but the value is increased to 90% using multiple conformations. After prediction of the complexes that are not available in the literature, new drug-target pairs are identified using a structurally clustered protein-protein interface dataset. Drugs that are suggested for repurposing are determined according to their binding free energy prediction via docking (Table 1) for the similar interfaces. The protein chain having a favorable binding energy for target-ligand complex is stated as the new target.

Three of the results involve EGFR-ERBB3 protein interface formed by structures with PDB IDs of 4UIP and 4LEO with chain IDs of A and C, respectively. EGFR is a transmembrane protein of ErbB family of tyrosine kinase receptors ^46^. EGFR, also known as HER1, involves extracellular region comprising four domains, transmembrane region and intracellular region with tyrosine kinase domain ^47, 48^. Domain III of the extracellular region plays a role in ligand binding ^46^. ERBB3, also known as HER3, is also a member of ErbB family and consists of three regions namely extracellular, transmembrane and intracellular regions. Its extracellular region also has four domains among which domains I and III are involved in binding of its natural ligand heregulin ^49^. EGFR and ERBB3 can form heterodimers as well as homodimers resulting in activation of MAPK/ERK and PI3K/Akt signaling pathways that are responsible for cell migration and proliferation ^50–52^. Previous studies showed that EGFR-ERBB3 heterodimer is involved in signaling which promotes metastasis in melanoma cells and activation of MAPK ^53^. According to another study, upregulation, mutation or catalytic activation of ErbB family proteins are associated with breast, ovarian, colorectal, pancreatic and lung cancer. Moreover, targeting a single protein in therapy might fail because of the crosstalk between ErbB family that activates downstream pathways. In that study, it is also reported that targeting the EGFR-ERBB3 interface for breast cancer is an improved strategy where malignancies exhibit resistance to treatment that targets a single protein ^54^.

Structures with PDB IDs of 4LEO and 4UIP are extracellular domains of ERBB3 and EGFR, respectively. A previous study suggested that targeting extracellular domain of EGFR is promising in colorectal cancer treatment where there is resistance to EGFR inhibitors cetuximab and panitumumab ^55, 56^. In our results, tipranavir and indinavir bind to EGFR with favorable binding energy at the interface formed between EGFR and ERBB3 (Figure 6). Both indinavir and tipranavir are drugs used in treatment of HIV infection ^57^. Tipranavir and indinavir were approved by FDA in 2005 and 1996, respectively ^58, 59^. They both bind to the active site of HIV protease enzyme to prevent hydrolysis of peptide bonds which is necessary for the life cycle of HIV ^60^. According to docking results, tipranavir and indinavir are bound to domain III of EGFR extracellular domain. In another study, cetuximab which is an EGFR inhibitor is also bound to the domain III ^61^ suggesting that these HIV protease inhibitors might be used in cancer treatment.

**Figure 6.**
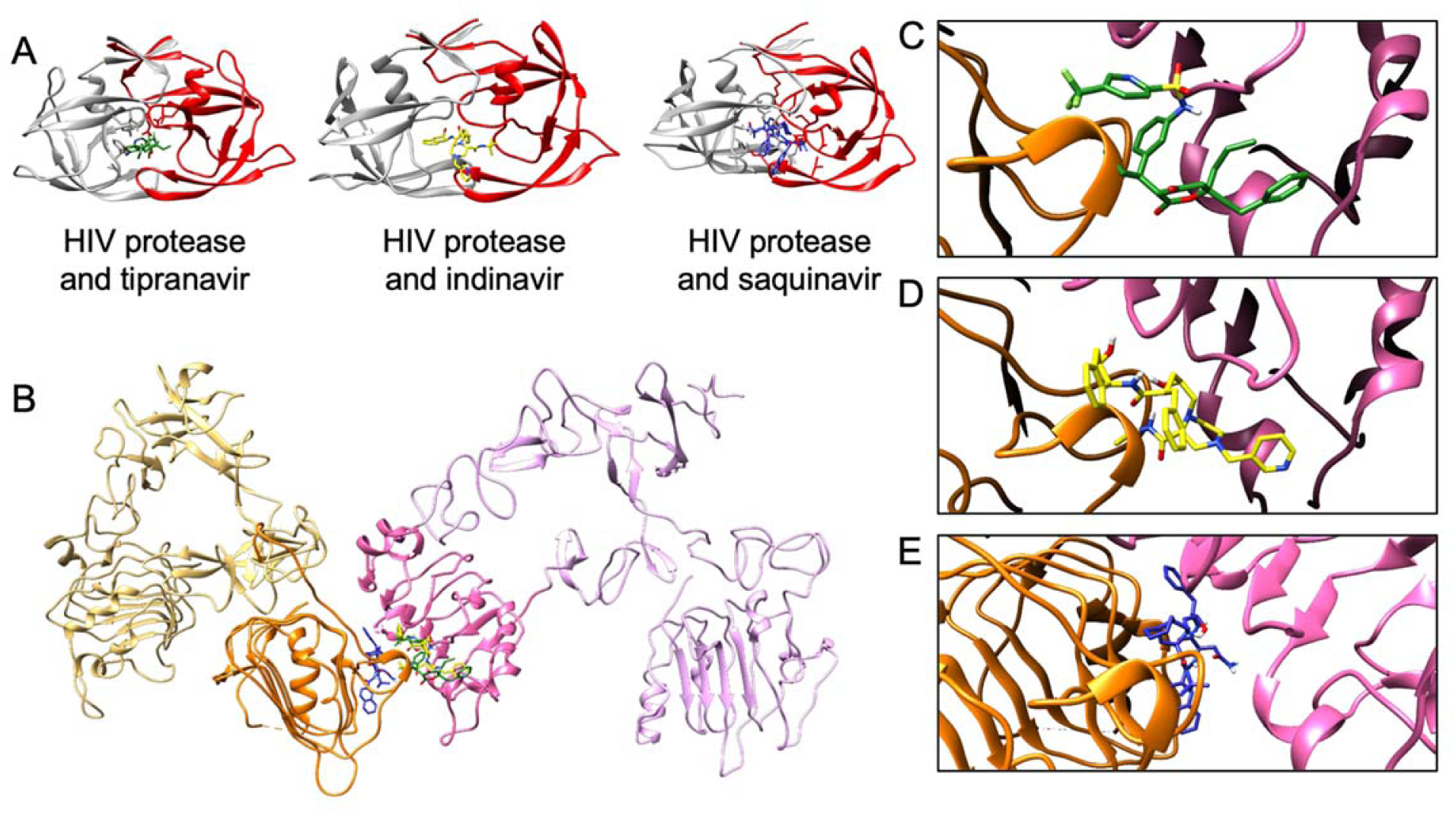
ERBB3-EGFR interface results. (A) The protein-protein interfaces that are in the same cluster as the template used by PRISM (2nxmAB) with tipranavir (1D4S_A_B), indinavir (1HSH_A_B) and saquinavir (1HXB_A_B). (B) ERBB3 (PDB ID:4LEO Chain ID:C), represented in orange with domain I in darker shade, and EGFR (PDB ID:4UIP Chain ID:A), represented in pink with domain III in darker shade ^32, 62^, in complex with tipranavir (forest green), indinavir (yellow) and saquinavir (medium blue). (C) Close-up of tipranavir bound to ERBB3-EGFR interface. (D) Close-up of indinavir bound to ERBB3-EGFR interface. (E) Close-up of saquinavir bound to ERBB3-EGFR interface.

The other drug that binds to the same interface formed by these protein structures is saquinavir which is also an HIV protease inhibitor. Saquinavir binds to ERBB3 with a lower (better) energy. Saquinavir was approved in 1995, being the first HIV protease inhibitor approved by FDA ^63^. Saquinavir is bound to domain I of ERBB3 extracellular domain, which is one of the domains that is involved in ligand binding and inhibition may prevent activation of downstream signaling pathways that play a role in growth of cancer cells.

Tipranavir and indinavir also bind to the interface that is formed between EGFR and ERBB2, with a lower binding energy to ERBB2 protein chain (Figure 7). The complex consists of chain A of structure with PDB ID 1YY9 and chain A of structure with PDB ID 3N85 representing EGFR extracellular domain and ERBB2 extracellular domain, respectively. ERBB2, also known as HER2, is another member of ErbB family. Thus, its extracellular domain consists of four subdomains where subdomains I and III are involved in ligand binding ^64^ and subdomains II and IV play roles in homodimerization and heterodimerization ^65^. EGFR-ERBB2 heterodimer activates MAPK pathway, preventing apoptosis ^66^. Overexpression of ERBB2 is highly related to breast cancer and is observed in 20-30% of all breast cancers ^67^. Upregulation of ERBB2 expression may promote cell proliferation and can further lead to tumorigenesis ^68^. Amplification of ERBB2 also occurs in 10-30% of gastric cancers and has been associated with different types of cancer such as ovary, colon and bladder cancers ^69–71^. Consequently, ERBB2 has become a therapeutic target of interest. Trastuzumab is a monoclonal antibody used in breast and gastric cancer and targets ERBB2 ^72, 73^. There are also other therapeutic strategies that are developed for patients with trastuzumab resistance. Dual tyrosine kinase inhibitor lapatinib is one of them and it targets both EGFR and ERBB2 ^74^. Moreover, recombinant humanized ERBB2 monoclonal antibody pertuzumab prevents dimerization of ERBB2 with EGFR and ERBB3 to prevent activation of downstream pathways which is demonstrated to be inhibiting breast and prostate tumor growth ^73, 75^. Since tipranavir and indinavir bind to the domain III of ERBB2 extracellular domain and is at the EGFR-ERBB2 interface according to our results, they may disrupt the heterodimer and prevent the cell signaling. Therefore, these HIV protease inhibitors may be repurposed for tumor growth inhibition. There have been studies on repurposing of HIV protease inhibitor nelfinavir for cancer, suggesting its mechanism of action involves inhibition of MAPK signaling pathway ^76, 77^. Moreover, phase II clinical trial of indinavir for non-HIV associated classic Kaposi’s Sarcoma, which is a cancer type, reported positive outcome after receiving treatment for 61.5% of the patients ^78^. Furthermore, a study demonstrated that tipranavir induced apoptosis of gastric cancer stem cells by targeting PRSS23-IL24 pathway ^79^. Hence, the HIV protease inhibitors that we reported in our results may be repurposing candidates for cancer.

**Figure 7.**
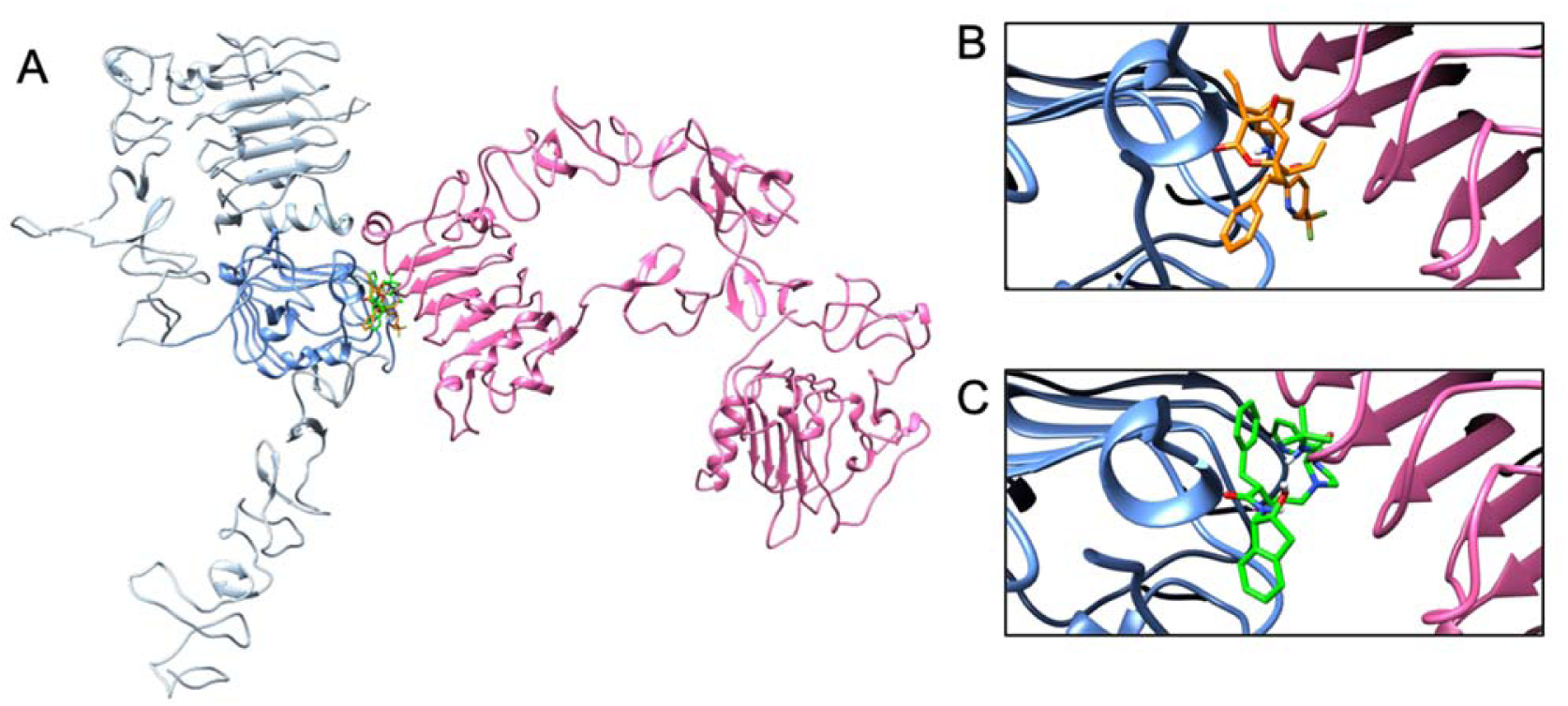
ERBB2-EGFR interface results. (A) ERBB2 (PDB ID:3N85 Chain ID:A), represented in blue with domain III in darker shade ^32, 62^, and EGFR (PDB ID:1YY9 Chain ID:A), represented in pink in complex with tipranavir (orange) and indinavir (green). (B) Close-up of tipranavir bound to ERBB2-EGFR interface. (C) Close-up of indinavir bound to ERBB2-EGFR interface.

The interface formed between RAF1 and BRAF also has drugs that have low binding energy. RAF1, also known as CRAF, and BRAF are both members of Raf kinase family along with ARAF. Their structure is comprised of three conserved regions (CR), namely, C1 with Ras-binding domain and cysteine-rich domain; CR2 with serine/threonine rich region; and CR3 involving kinase domain ^80^. Heterodimer of BRAF and RAF1 formation is induced by growth factor-stimulated RAS and activates MEK and ERK to promote cell proliferation, differentiation, survival and migration ^81, 82^. BRAF-RAF1 heterodimer is the most active dimer compared to their homodimers in MEK1/2 activation ^83, 84^. BRAF mutation is observed in nearly 8% of all cancers and mostly associated with melanoma ^85^. Mutated RAF1 is less common in human cancers but mutation in RAF1 may lead to Noonan syndrome which is a disorder that includes short stature, facial dysmorphology and congenital heart defects ^86, 87^. Also, it is reported that increased BRAF heterodimerization with RAF1 is associated with RAF1 mutations related to Noonan syndrome ^88^. Since mutation in BRAF also promotes MAPK signaling pathway activation and consequently tumorigenesis, it has been identified as a target in cancer therapy ^83^.

According to our results, granisetron and galantamine bind to BRAF with favorable energy at the interface formed between BRAF (PDB ID:6Q0K Chain ID:A) and Ras binding domain and cysteine-rich domain of RAF1 (PDB ID:6XGU Chain ID:B) (Figure 8). Granisetron is a serotonin type 3 (5-HT3) receptor antagonist used as antinauseant for cancer chemotherapy patients ^89^. There are several studies where some other drugs binding to a serotonin receptor are proposed as anticancer agents. For example, tegaserod which is a serotonin receptor 4 (HTR4) agonist is reported to be inducing apoptosis in B16F10 murine melanoma cell line and some human melanoma cell lines by perturbing PI3K/Akt/mTOR pathway ^90^. In another study, methiothepin which is a nonselective serotonin 5-HT receptor antagonist is reported to be increasing the efficacy of chemotherapy when used along with doxorubicin, against melanoma cells ^91^. In the same study, it is shown that methiothepin also enhances the efficacy of BRAF inhibitor vemurafenib and MEK inhibitor trametinib, used against resistant BRAFV600E melanoma cells.

**Figure 8.**
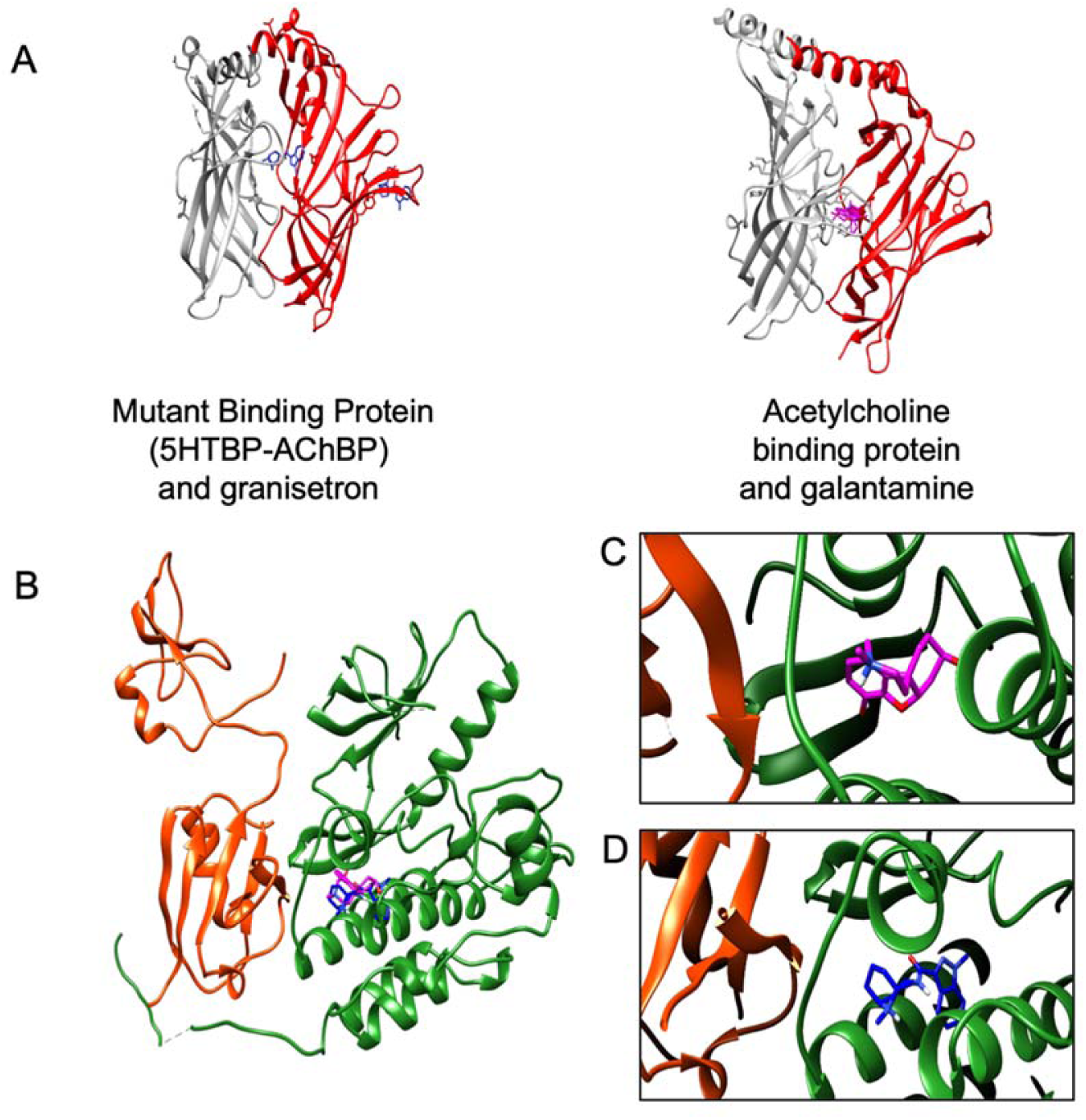
BRAF-RAF1 interface results. (A) The protein interfaces that are in the same cluster as the template used by PRISM (1uv6GH) with granisetron (2YME_A_B) and galantamine (2PH9_C_D). (B) BRAF (PDB ID:6Q0K Chain ID:A), represented in green and RAF1 (PDB ID:6XGU Chain ID:B), represented in orange in complex with granisetron (medium blue) and galantamine (magenta). (C) Close-up of galantamine bound to BRAF-RAF1 interface. (D) Close-up of granisetron bound to BRAF-RAF1 interface.

Galantamine is an acetylcholinesterase inhibitor used in treatment of Alzheimer’s disease ^92^. Abnormal expression of acetylcholinesterase is observed in several tumors, therefore, is associated with tumor development ^93–98^. As a result, some acetylcholinesterase inhibitors may be considered as possible anti-cancer agents for the cancer types where increased activity of acetylcholinesterase is observed ^99^. Inhibition of the MAPK pathway may be another mechanism when using acetylcholinesterase inhibitor galantamine as an anti-cancer agent.

Both granisetron and galantamine are bound to the kinase domain ^100, 101^ according to our results. BRAF inhibitors such as sorafenib also binds to kinase domain of BRAF (PDB ID:1UWH) and if these drugs also act as BRAF inhibitors or disrupt the BRAF-RAF1 protein interface, they can be potential anti-cancer drugs. However, in some cases, a BRAF inhibitor such as vemurafenib, binding to BRAF leads to inhibition of BRAF but transactivation of RAF1 further leads to activation of MEK and ERK. To prevent paradoxical activation, a high level of RAF inhibitor that acts on both BRAF and RAF1 may be used ^102^.

Additionally, somatic mutations in human cancers are mapped to interfaces of the 3-dimensional structures of the protein complexes used in this study via COSMIC database ^103^ and SIFTS UniProt-PDB mappings ^104^ on PDBe API ^105^. COSMIC database provides manually curated mutation information of tumor samples including mutation types. Here, nonsense mutations that stop the translation prematurely and missense mutations that result in encoding of different amino acid at that location are mapped. For the ERBB3-EGFR complex, the interface mutation with the highest frequency for ERBB3 is observed in 0.006% of the samples and they are from endometrium, large intestine and bile duct tumors whereas the highest frequency for the interface mutations of EGFR is 0.007%, from the samples of large intestine and lung carcinoma (Table S9). For the BRAF-RAF1 complex, the highest frequency of mapped mutations is 0.005% for RAF1 and they are from various tissues such as large intestine, brain and endometrium. For BRAF, it is 0.003% from mutations in the samples of ovary, lung, kidney, skin and lymphoid. At the interface of ERBB2-EGFR complex, ERBB2 has mutations with the frequency of 0.002% observed in tissues like skin, ovary and stomach whereas the highest mutation frequency for EGFR is 0.007%, observed in large intestine and lung tissues. The protein structures that we used in our studies don’t exhibit these mutations and the mutations may change the protein-protein interactions and the interaction with the drug. However, not all of the people with cancer have these mutations and considering the frequency of the mutations among the tumor samples, the frequency is not very high.

This study relying on structural similarities of protein-protein interfaces revealed that indinavir, tipranavir and saquinavir originally used for HIV infection treatment may bind to EGFR-ERBB3 and/or EGFR-ERBB2 interfaces and can be repurposed for cancer treatment. Additionally, Alzheimer’s disease drug galantamine and antiemetic drug granisetron may bind to BRAF-RAF1 interface and can be used as anti-cancer agents to prevent tumor growth. Even though these results present candidates for drug repurposing, they should be validated by experiments and clinical trials.

## CONCLUSIONS

Drug repurposing is a strategy that can be adopted to save time and money by reducing drug development timeline and research and development process cost. Hence, it is an effective alternative to conventional drug development. There are different approaches for drug repurposing involving methods based on similarities in drugs, targets or diseases. Here, we focused on the structural target similarities considering protein-protein interfaces formed by proteins involved in Ras/Raf/MEK/ERK signaling pathway. This pathway plays role in cell signaling that regulates cell proliferation, differentiation, apoptosis; therefore, is highly related to cancer and tumor progression. The protein-protein interfaces studied in this work either have been predicted by PRISM according to physically interacting proteins in STRING database or obtained from Protein Data Bank. Candidates for drug repurposing are suggested considering the binding free energy prediction of the drug to the protein interface that is structurally similar to its original target by docking.

We report that HIV protease inhibitors tipranavir, indinavir and saquinavir can bind to EGFR-ERBB3 interface. Additionally, tipranavir and indinavir can bind to EGFR-ERBB2 interface. Furthermore, we report that galantamine used in Alzheimer’s disease treatment and antiemetic drug gransiteron can bind to RAF1-BRAF interface. These protein interfaces are involved in signal transduction that activates Ras/Raf/MEK/ERK signaling pathway leading to biological processes that promote tumor growth. Hence, disruption of these interfaces may interrupt transduction of the signals associated with cancer. Consequently, these drugs are proposed to be repurposed as anti-cancer agents.

Although our results present some candidates for drug repurposing and are important in identification of the compound to be repurposed, in-silico drug repurposing approach needs to be supported by experimental data that shows complete effect of the drug. Thus, candidates suggested in this work should be validated experimentally and by clinical trials in the future studies.

## MATERIALS AND METHODS

The basis of this study is that if a drug can bind to a protein-protein interface, it may also bind to another interface which is structurally similar to the protein-protein interface that the drug is originally bound to. Since our study focuses on Ras/Raf/MEK/ERK signaling pathway, we extracted the structures of the proteins in this pathway. Then, the alternative conformations of these proteins are determined and used in the prediction of the complexes of physically interacting proteins if they are not available in the literature. Following that, protein-protein interfaces with drugs are filtered to suggest new targets to these drugs using a structurally similar protein-protein interface dataset and docking. Figure 9 illustrates the workflow of this study.

**Figure 9.**
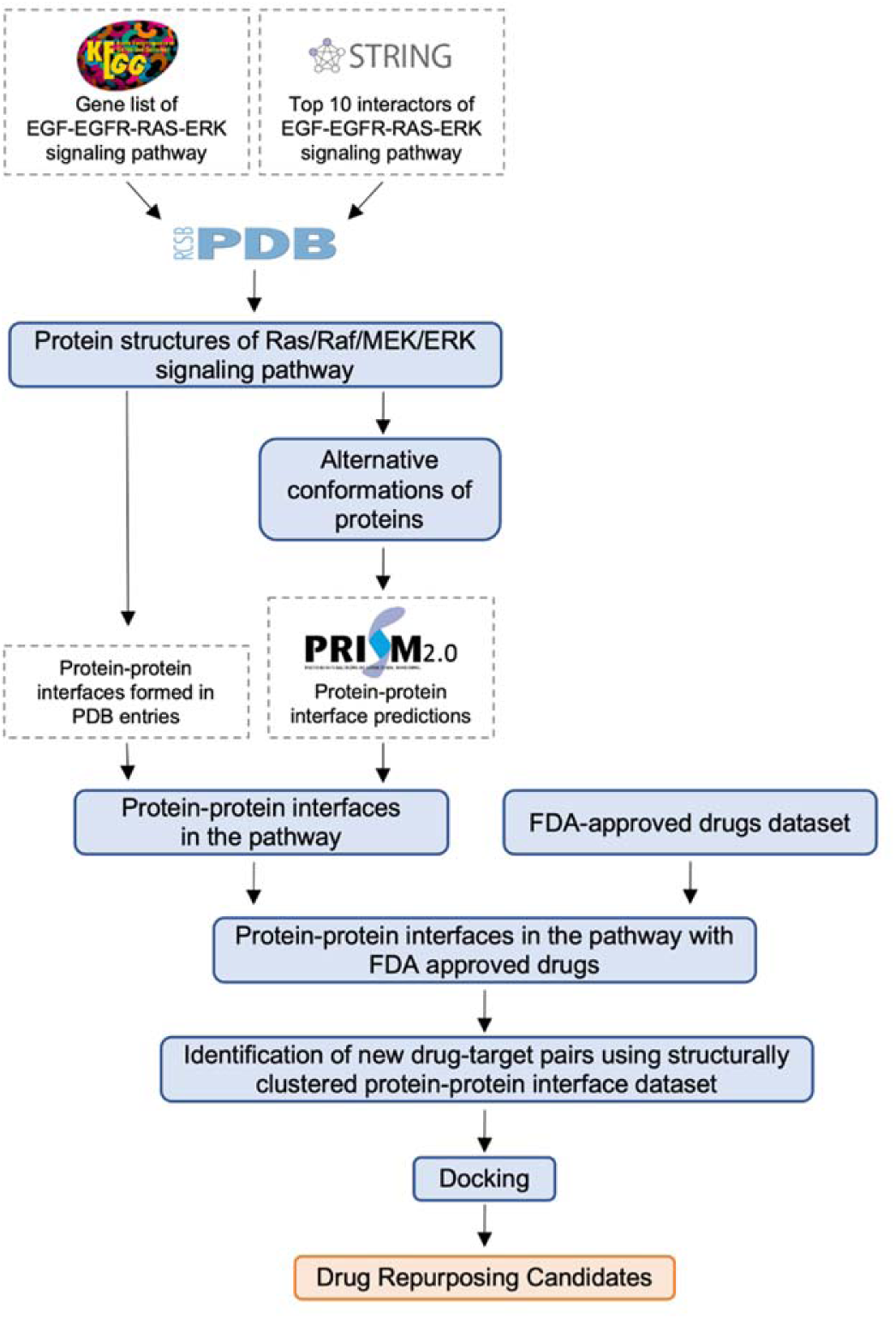
Workflow of the study. A list of proteins in the Ras/Raf/MEK/ERK signaling pathway is constructed from the KEGG and STRING databases. Their three-dimensional structures are found in the Protein Data Bank (PDB). These structures are grouped to obtain alternative conformations. The protein-protein interfaces formed by these proteins in PDB entries are determined or predicted via a template-based protein-protein docking tool. The interfaces with approved drugs are filtered using FDA-approved drugs dataset and new drug-target pairs are identified using a structurally clustered protein-protein interface dataset. Then, docking is performed to propose drug repurposing candidates.

### Protein structures of Ras/Raf/MEK/ERK signaling pathway

The gene list for the EGF-EGFR-RAS-ERK signaling pathway (N00001) under MAPK signaling pathway is obtained from the Kyoto Encyclopedia of Genes and Genomes (KEGG) ^36–38^. KEGG identifiers for these genes are mapped to UniProt identifiers. If there are more than one UniProt identifier associated with the gene, the UniProtKB/Swiss-Prot identifier (reviewed, manually annotated) is selected. Physical interactions for the 16 proteins in the EGF-EGFR-RAS-ERK signaling pathway with highest confidence score (≥0.900) and their top 10 interactors (Table S10) are imported from the STRING database ^39^. The proteins in EGF-EGFR-RAS-ERK signaling pathway and their top 10 interactors form our set of pathway proteins.

Following that, PDB entries for these UniProt identifiers are found using “idmapping_selected.tab.gz” file from the UniProt website ^106^. Since PDB is redundant and some PDB entries are very similar, proteins with 95% sequence identity and 2 Å RMSD value are grouped for each UniProt identifier. One representative is kept for each group. Proteins having less than 30 residues are eliminated ^107^. These steps provided us with multiple conformations of the pathway proteins introducing dynamics in the predictions.

### Protein-protein interfaces in the pathway

Protein-protein interfaces used in this work are either predicted by the PRISM web server or found in PDB. PRISM predicts interactions between two proteins according to the similarity between the surfaces of target proteins and each side of a template interface. Physically interacting protein pairs according to STRING are sent to the PRISM web server as target proteins. PRISM results consist of an interface template, binding energy, protein complex structure and interface residues. PRISM may give none or multiple results for each target protein pair.

Additionally, PDB entries involving one of the proteins in the EGF-EGFR-RAS-ERK signaling pathway are found. Only the proteins listed under the EGF-EGFR-RAS-ERK signaling pathway in KEGG are included to avoid 2^nd^ shell interactors. Protein-protein interfaces formed in these PDB entries are used in the following steps. If the distance between two atoms is less than the sum of their van der Waals radii plus a tolerance of 0.5 Å, those are considered contacting. If there are at least five contacting residues at each protein chain, they are considered to be forming an interface.

### Filtering of protein-protein interfaces with Food and Drug Administration (FDA) approved drugs

FDA-approved drugs are listed in the ZINC database ^108^ and those in PDB are identified to form FDA-approved drugs dataset used in this work (Dataset S1). Glycerol (PDB Ligand ID:GOL) and isopropyl alcohol (PDB Ligand ID:IPA) that are present in FDA-approved drugs dataset are highly observed in PDB entries. However, these molecules are mostly used in the structure determination step as precipitant or to protect proteins when frozen ^109, 110^. Hence, glycerol and isopropyl alcohol are excluded from FDA-approved drugs dataset in this step.

Protein-protein interfaces in the EGF-EGFR-RAS-ERK signaling pathway with FDA-approved drugs are filtered by mapping and combining ligands at the interface residues using data from PDBsum ^111^ with the FDA-approved drug dataset (Dataset S2). Protein-protein interfaces predicted by PRISM and interfaces from the PDB are studied separately.

### Identification of new drug-target pairs

To propose new drug target pairs, a dataset consisting of clusters of structurally similar protein-protein interfaces is used (Dataset S3) ^43^. This dataset is constructed by clustering protein interfaces in PDB entries with an Interface-Similarity score (IS-score) of 0.311 according to iAlign ^112^. SparseHC ^113^, a hierarchical clustering algorithm, is used in the clustering.

Two cases are considered to propose new drug-target pairs (Figure 3):

Repurposing To: A drug bound to one of the interfaces in a protein-protein interface cluster may bind to an interface in the same cluster, and the protein is in the Ras/Raf/MEK/ERK pathway.

Repurposing From: A drug bound to a protein interface in Ras/Raf/MEK/ERK pathway may also bind to another protein interface that is in the same cluster, and the protein is not in the Ras/Raf/MEK/ERK pathway.

### Docking

Python package of AutoDock Vina ^114^ is used in this work for docking. Additionally BioPython ^115^ and NumPy packages ^116^ in Python, Chimera ^117^ and Open Babel ^118^ are used. 3-dimensional structure of drugs at reference pH are downloaded from ZINC database ^108^. Both receptor and ligand structures are prepared for docking using codes in MGL Tools ^119^. The size and the center of the docking box is adjusted to include interface residues (Dataset S4) in the box. Docking is performed with exhaustiveness of 8 because it is the best option for prediction of binding energy considering the increased computation time with a higher exhaustiveness ^44^. Details are presented in Text S2.

### Cancer Mutations

For the somatic mutations in cancer, missense and nonsense mutations from the COSMIC (v97) database are used ^103^. Human proteins included in the proteome UP000005640 ^106^ are considered. Using PDBe API ^105^, UniProt mappings for each PDB ID from SIFTS ^104^ are obtained. For the start and end residue numbers, author residue numbers are considered. The residue names are compared, the start and end residues are manually adjusted to be consistent if they don’t match. The mutations in the whole chain are listed and compared with interface residues to find the ones that are at the interface.

## DATA AND SOFTWARE AVAILABILITY

PRISM is accessible through the PRISM webserver (https://cosbi.ku.edu.tr/prism/). The codes forgrouping of the alternative conformations are available on GitHub (https://github.com/ku-cosbi/ppi-network-alternative). All PRISM results and docking results of the proposed drugs are available on GitHub (https://github.com/ku-cosbi/RasPathwayDrugRepurposing). Other data used in this work can be found in supplementary materials.

## SUPPORTING INFORMATION

**Table S1. Group Representatives of Ras/Raf/MEK/ERK Pathway Protein Structures in PDB.** The representatives of the alternative conformation groups of the pathway proteins (XLSX).

**Table S2. Possible drug-target pairs for “Repurposing To”.** Identified drug target pairs with “Repurposing To” approach (XLSX).

**Table S3. FDA Approved Drugs at Protein Interfaces Formed by Ras/Raf/MEK/ERK Pathway Proteins.** List of drugs that are contacting to interface residues of protein-protein complexes in Ras/Raf/MEK/ERK pathway (XLSX).

**Table S4. Possible drug-target pairs for “Repurposing From”.** Identified drug target pairs with “Repurposing From” approach (XLSX).

**Table S5. All docking binding free energy results.** Binding free energies of the drug to the both sides of the protein-protein interface (XLSX).

**Table S6. Drug repurposing candidates detailed results.** PRISM complex name and all the interfaces with the drug that is structurally similar are presented (XLSX).

**Table S7. Cancer mutations at the protein-protein interfaces for “PRISM interfaces”.** Cancer mutations at the interface residues for the interfaces predicted by PRISM (XLSX).

**Table S8. Cancer mutations at the protein-protein interfaces for “Interfaces from PDB”.** Cancer mutations at the interface residues for the interfaces that are already available in PDB and formed by the pathway proteins (XLSX).

**Table S9. Frequency and tissue information for interface mutations.** Detailed frequency, tissue and histology information for the interface mutations mapped from COSMIC to ERBB3-EGFR, BRAF-RAF1 and ERBB2-EGFR complexes (XLSX).

**Table S10. Physical interactions of pathway proteins.** List of physically interacting protein pairs (XLSX).

**Text S1. PRISM results.** Protein complexes predicted via PRISM (XLSX).

**Text S2. Details of docking.** More detailed information on the docking procedure followed in this work is presented (XLSX).

**Dataset S1. List of FDA approved drugs.** Set of FDA approved drugs that are on PDB (XLSX).

**Dataset S2. Protein-protein interfaces with FDA approved drugs.** Dataset of the protein-protein interfaces with FDA approved drugs that are on PDB and used in this work (XLSX).

**Dataset S3. Structurally clustered protein-protein interface dataset.** Each line represents a cluster (XLSX).

**Dataset S4. Contacting residues of the protein-protein interfaces used in this work.** Each line represents an interface (XLSX).

## ACKNOWLEDGEMENT

We would like to thank Prof. Ruth Nussinov and Dr. Hyunbum Jang for their valuable comments. This project has been partially funded by TUSEB 4448/4081 and TUBITAK 2247-120C120.

